# *Phenomenal*: An automatic open source library for 3D shoot architecture reconstruction and analysis for image-based plant phenotyping

**DOI:** 10.1101/805739

**Authors:** Simon Artzet, Tsu-Wei Chen, Jérôme Chopard, Nicolas Brichet, Michael Mielewczik, Sarah Cohen-Boulakia, Llorenç Cabrera-Bosquet, François Tardieu, Christian Fournier, Christophe Pradal

**Author notes:** Co-corresponding authors: Christian Fournier, Christophe Pradal. **Note**: The last two authors contributed equally to the leading of this work. **Author contributions**: S.A. developed the software with contributions of J.C., N.B., M.M., C.F., and C.P.; F.T., C.F. and C.P. conceived the research plans; T.W.C and L.C.B. performed the PhenoArch experiments with contributions of N.B., C.F., and S.A.; T.W.C., M.M., S.A., C.F. analysed the data; S.C.B and C.P. conceived and designed the software and big data infrastructure; S.A, T.W.C, F.T, C.P., C.F. wrote the article with contributions of all authors F.T, C.F. and C.P. supervised and completed the writing; C.F and C.P. agree to serve as the author responsible for contact and ensure communication. **One-Sentence summary:** Automatic 3D shoot reconstruction and analysis from phenotyping images.

## Abstract

In the era of high-throughput visual plant phenotyping, it is crucial to design fully automated and flexible workflows able to derive quantitative traits from plant images. Over the last years, several software supports the extraction of architectural features of shoot systems. Yet currently no end-to-end systems are able to extract both 3D shoot topology and geometry of plants automatically from images on large datasets and a large range of species. In particular, these software essentially deal with dicotyledons, whose architecture is comparatively easier to analyze than monocotyledons. To tackle these challenges, we designed the *Phenomenal* software featured with: (i) a completely automatic workflow system including data import, reconstruction of 3D plant architecture for a range of species and quantitative measurements on the reconstructed plants; (ii) an open source library for the development and comparison of new algorithms to perform 3D shoot reconstruction and (iii) an integration framework to couple workflow outputs with existing models towards model-assisted phenotyping. *Phenomenal* analyzes a large variety of data sets and species from images of high-throughput phenotyping platform experiments to published data obtained in different conditions and provided in a different format. Phenomenal has been validated both on manual measurements and synthetic data simulated by 3D models. It has been also tested on other published datasets to reproduce a published semi-automatic reconstruction workflow in an automatic way. *Phenomenal* is available as an open-source software on a public repository.

## Introduction

Morphological traits determining 3D plant architecture have a large effect on biomass accumulation via light interception, light distribution in the canopy and light-use efficiency (Zhu et al., 2010, Chen et al., 2018; Bucksch et al., 2017; Balduzzi et al., 2017). *In silico* studies using dynamic 3D plant architecture suggest that biomass can be increased by up to 20% by optimizing architectural traits (Chen et al., 2014; 2015). Examining the genetic control of these traits is an avenue for crop improvement that has been under-exploited until now, but requires high throughput and reproducible phenotyping pipelines that are lacking until now (Tardieu et al., 2017). Several techniques are available for capturing 3D plant morphology and geometry, *e.g.* manual plant digitizing (Godin et al., 1999; Danjon et al., 2008; Vos et al., 2010), stereoscopic plant modelling (Biskup et al. 2007; Lati et al. 2013), laser scanning (Paulus et al., 2014), multi-view image reconstruction (Pound et al., 2014; Nguyen et al., 2017; Gibbs et al., 2017, 2018; Pfeifer et al., 2018) or depth imaging (McCormick et al., 2017). However, most of them have not been ported to high throughput. Phenotyping platforms allow fulfilling this need via rapid and automatic collection of multi-view 2D above-ground plant images for thousands of plants (Fiorani and Schurr, 2013; Tardieu et al., 2017).

Most existing pipelines approach plant architecture via a series of independent 2D analyses (e.g., Knecht et al., 2016; Zhang et al., 2017) using regressions between traits of interest and 2D image features such as pixel counts, convex hull area or connectivity (Walter et al., 2007; Jansen et al., 2009; Hartmann et al., 2011; Klukas et al., 2014; Knecht et al., 2016; Burgess et al., 2017; Gibbs et al., 2018). Recently, Das Choudhury et al. (2018) managed to extract morphological traits such as leaf length, leaf angle and leaf area, using a pipeline based on 2D skeleton analysis of maize images. However, 2D skeletons have intrinsic limitations due to 2D projection, such as angular deformation and crossing artifacts. As a consequence, the number of biological traits that can be quantitatively and automatically derived from image processing is still limited and does not approach the potential of 3D plant imaging.

The few automatic methods capturing plant 3D features are specific to one species or category of species (Gibbs et al., 2017). Most pipelines are limited to dicotyledonous plants (Paproki et al., 2012), for which leaf segmentation is simplified by easily identifiable petioles. Elnashef et al. (2019) showed that tensor-based classification of 3D point cloud allowed stem detection on maize, wheat and cotton. Still, the leaf segmentation task remains complex in monocotyledonous species where leaves are directly connected to the stem, often rolled within each other and frequently crossing neighboring leaves (Das Choudhury et al., 2018, Elnashef et al., 2019).

Currently, existing 3D analysis pipelines are either semi-automatic (Mccormick et al., 2017), automatic but only for a very limited dataset (Elnashef et al., 2019), or implement only a subpart of the phenotyping pipeline, like 3D geometric reconstruction or organ segmentation from a mesh (Paproki et al., 2012; Pound et al., 2014). As a consequence, to our knowledge, none of these 3D solutions has been tested on large datasets.

We present here the open source library *Phenomenal* for building automated high throughput analysis pipelines such as 3D shoot reconstruction, light interception, organ segmentation, or complex trait extraction (https://github.com/openalea/phenomenal). *Phenomenal* pipelines are fully compatible with the scientific workflow infrastructure InfraPhenoGrid (Pradal et al., 2017) that allows processing of large dataset on the cloud (Heidsieck et al., 2019), and with the open-source OpenAlea scientific workflow system that provide full provenance of computations and ensure their reproducibility (Cohen-Boulakia et al., 2017) (https://github.com/openalea/openalea). Here, *Phenomenal* was first used to build a 3D shoot reconstruction pipeline for several species in 8 high-throughput experiments (Cabrera-Bosquet et al., 2016). Then, the automated architectural analysis pipeline using the novel organ segmentation algorithms developed in *Phenomenal* was challenged on maize and sorghum using three independent tests.

## Results

### A library for automatic 3D topology and geometry reconstruction from images

The *Phenomenal* library provides automatic, integrated and robust image analysis pipelines (Fig. 1, ABCD) for reconstructing 3D topology and geometry of plants on high-throughput multi-view images, based on (i) a camera calibration procedure adapted for multiple fixed-cameras with a rotating object system (Fig. 1.A), (ii) a robust background subtraction filter based on a mean shift algorithm (Brichet et al., 2017) (Fig.1.B), (iii) a 3D voxel volume reconstruction (Fig. 1.C) of the plant using an improved and fast space-carving algorithm (Kutulakos et al., 2000). To speed up the last algorithm, we use an octree decomposition technique (Slabaugh et al., 2001) to carve the 3D volume. We also improve the algorithm to better support common imperfections of high-throughput plant imaging systems (plant movements, light reflections, binarization errors) that may cause parts of the plant to disappear (Supplementary Material S1-Robust multi-view reconstruction). Then, the marching cube algorithm (Lorensen & Cline, 1987) transforms the 3D voxel volume in a standard 3D mesh (Fig. 1D).

**Figure 1.**
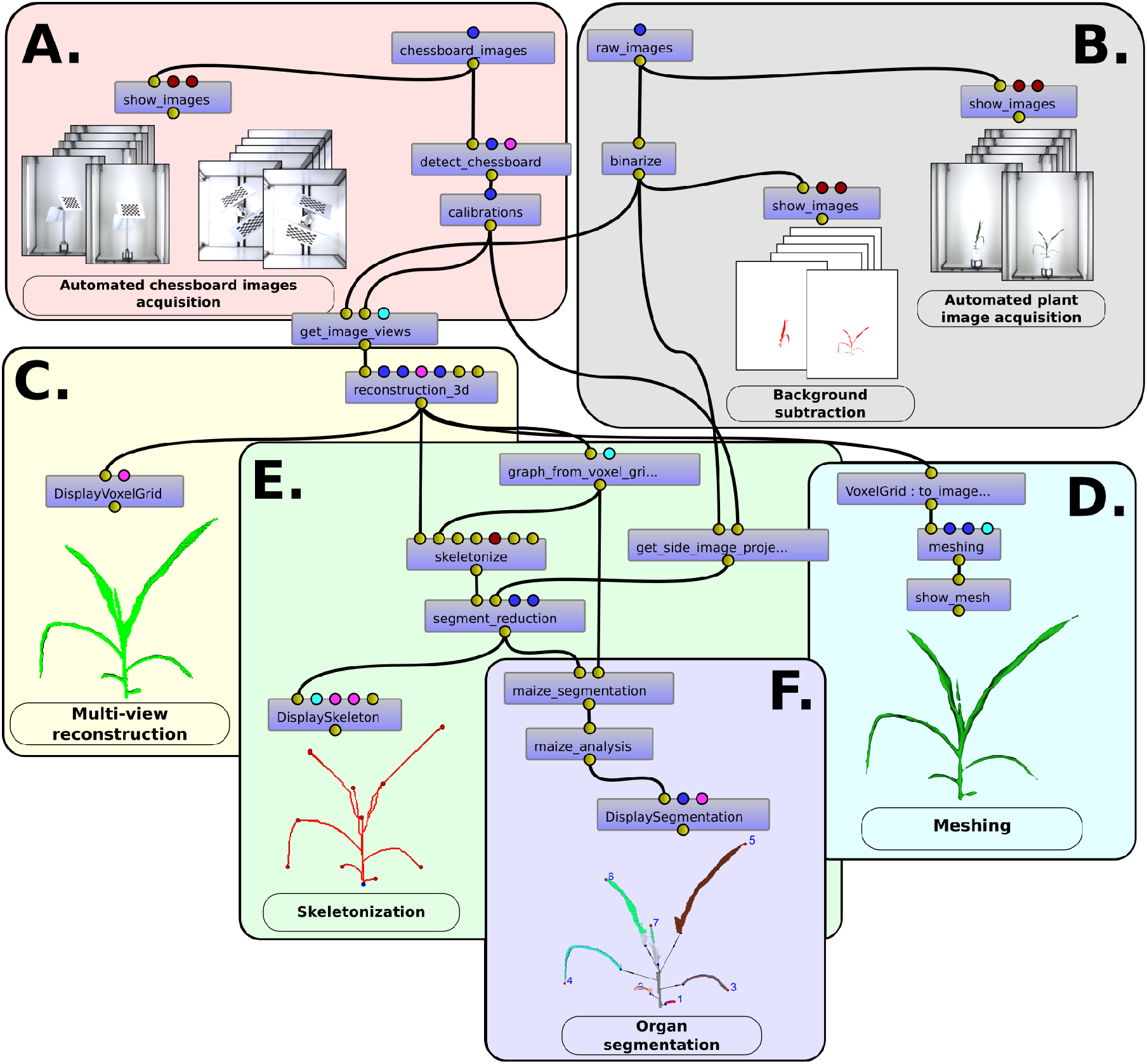
Snapshot of an image processing pipeline composed with *Phenomenal* under the visual programming environment VisuAlea of the OpenAlea platform, superposed with graphical outputs produced by terminal point the workflow. Small blue boxes are for atomic algorithm of the Phenomenal library. Small disc above and below boxes represent the inputs and outputs ports, the color of the disc being related to the data type. Connections between ports indicate which output is reused as an input for another algorithm. Large boxes delimit subparts of the workflow used to produce to the displayed output. (A) Multiple camera calibration before starting an experiment; (B) plant segmentation by background subtraction on the different views of the plant; (C) 3D geometric multi-view reconstruction from plant images; (D) 3D mesh production (E) skeleton computation from the 3D plant; (F) 3D organ segmentation and architectural trait analysis used for genome-wide association study.

Second, Phenomenal provides new algorithms, based on 3D skeleton analysis, to perform individual organ segmentation and morphological analysis of the plant. Because such segmentation is already available for dicotyledons based on planar / tubular segmentation techniques developed elsewhere (Paproki et al., 2012, Elnashelf et al., 2019), our algorithm also targeted monocotyledonous plants, where oblong leaves are directly attached to stems and are partly tubular. (Fig. 1. E and F). Standard skeletonization algorithms produced results that were too noisy, so a new algorithm was designed to identify a unique path per leaf that allows connecting the plant base with leaf tip (Supplementary Material S1-3D skeletonization).

Finally, these paths are used to segment the plant into stem and leaves, either expanding or mature (Supplementary Material S1 – Organ Segmentation). The final outputs of *Phenomenal analysis* workflows can be either a volume, a 3D-mesh, or a topology with its associated geometrical traits (Fig. 1F and Fig. 2).

**Figure 2.**
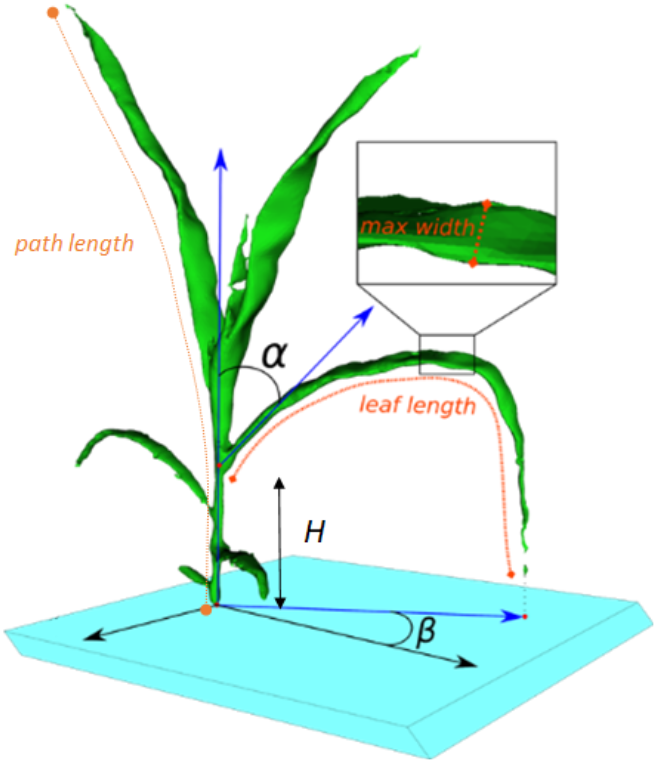
Architectural traits measured with the *Phenomenal* analysis algorithm. Measured traits displayed are stem height (*h*), path length, leaf length, average and maximal leaf width, leaf insertion angle (*α*) and azimuth angle (*β*)

### Phenomenal allowed automatic 3D reconstruction of 8 high throughput experiments with four different species

We tested 3D reconstruction analysis workflow by processing data of eight experiments (Table 1), in which 1680 plants were imaged daily for 30 to 40 days with either 3 or 13 images of each plant every day, obtained with two calibrated camera positioned on the side and the top of the plant. Each experiment involved up to 955,372 RGB images (Table 1) that are stored in the local instance of the Phenotyping Hybrid Information System (PHIS, Neveu et al., 2019).

**Table 1.** Summary of the workflow output from eight PhenoArch experiments. In experiments ZB12 and MA14, images of the plants were only acquired from two side views and one top view. Success rate (%) represents the amount of plants, which was successfully automatically binarized and reconstructed.

| Experiment (season) | Species | Number of binarized images | Number of reconstructed 3D plants | Success rate (%) | Number of views | Reference |
| --- | --- | --- | --- | --- | --- | --- |
| ZB12 (Spring) | Maize | 54 870 | 18 242 | 100 | 3 | Alvarez-Prado et al. (2017) |
| ZA13 (Winter) | Maize | 304 519 | 14 755 | 99.98 | 13 | Alvarez-Prado et al. (2017)<br>Chen et al. (2018) |
| ZB13 (Spring) | Maize | 237 782 | 12 940 | 99.95 | 13 | Alvarez-Prado et al. (2017)<br>Chen et al. (2018) |
| MA14 (Winter) | Apple | 64 317 | 21 439 | 99.98 | 3 | Lopez et al. (2015) |
| ZB14 (Fall) | Maize | 549 432 | 31 521 | 99.98 | 13 | Unpublished |
| GA16 (Spring) | Cotton | 639 405 | 47 990 | 99.95 | 13 | Unpublished |
| ZA16 (Spring) | Maize | 553 056 | 39 548 | 99.47 | 13 | Alvarez-Prado et al. (2017)<br>Chen et al. (2018), |
| ZA17 (Spring) | Maize | 955 372 | 73 385 | 99.41 | 13 | Brichet et al. (2018)<br>Perez et al. (2019) |

Three millions of images belonging to four species were requested from PHIS during the computational experiments, and processed automatically by the 3D reconstruction workflow (Table 1, Fig. 3). Regardless of the species (monocotyledon or dicotyledon), the success rate was higher than 99%, with only two experiments where the success was between 99 and 99.95% due to a technical disruption of the imaging system (Table 1). After curation of the database by tagging corrupted images and fixing inconsistencies, the success rate reached 100% for all experiments. Only the binarization parameters (threshold values) had to be slightly modified between experiments to account for changes in image acquisition settings (camera or cabin replacements). The quality of the reconstructed plant volume was sensitive to the number of views, as demonstrated on maize with larger (nosier) reconstructed volume in an experiment (ZB12) involving 3 views per plant instead of 13 for other maize experiments.

**Figure 3.**
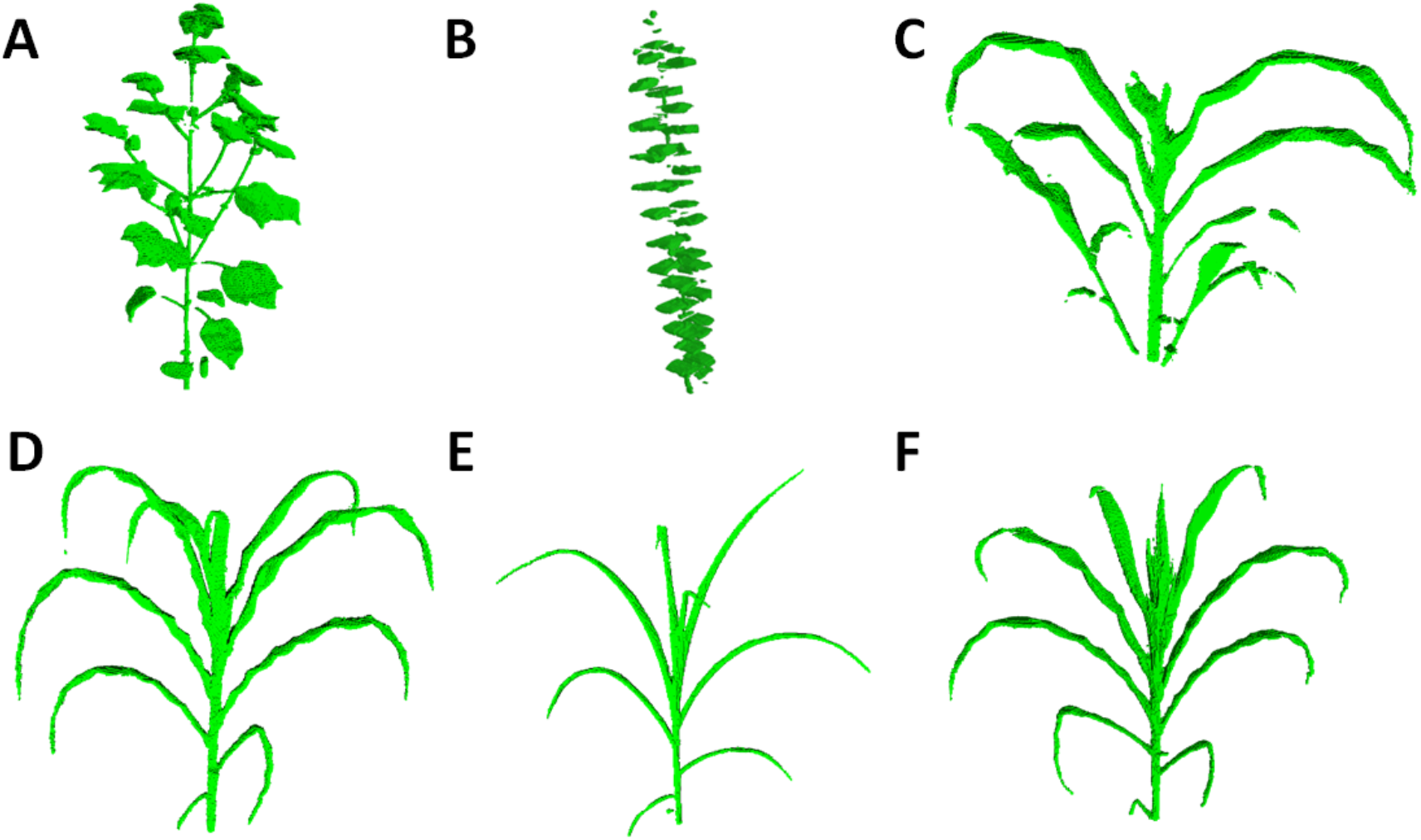
3D reconstruction of different plant species imaged in the PhenoArch platform with *Phenomenal*: (A) Cotton (*Gossypium*); (B) Apple tree (*Malus pumila*); (C) Sorghum (*Sorghum bicolor*); (D-F) Three Maize (*Zea Mays L.*) genotypes (EZ38, A375 and PHZ51) with contrasting plant architecture

We first tested a large diversity of maize genotype (342 lines and 402 hybrids), a large number of stages of development (i.e. from seedlings to mature plants), and well watered and water-stressed plants. The quality of reconstruction was satisfactory in all cases. The limited number of viewpoints (13), aimed at keeping a high throughput, and the undulation of leaf margin in this specie resulted however in reconstructed leaves being thicker than in reality. A loss of details can also be noticed in the whorl region (e.g. Fig. 3F), where small growing leaves tightly rolled within larger leaves are at best reconstructed as small tips emerging from a compact englobing volume. Sorghum plants (Fig. 3C), that display an architecture similar to maize but with lateral ramifications, were equally satisfactory reconstructed with 13 views, although some occlusions may produce some defects near plant base. Young Apple Trees reconstructed with 3 views (Fig 3B) allowed to clearly distinguish all leaves along the stem, but probably lack of supplemental viewpoints to capture with details individual leaves. Finally, the experiment on Cotton with 13 views (Fig. 3A) demonstrate the ability to reconstruct a very different kind of architecture mixing small thin structure (petioles) and large flat leaves.

### Phenomenal was successfully tested against manual measurements, published datasets and synthetic datasets

The algorithm was tested on three independent dataset (Fig. 4). The first test consisted of comparing outputs of the pipeline with manual measurements performed during one high-throughput experiment on *PhenoArch (*Fig. 4 A, B*, and* C*; Supplementary Material S2)*. A second test challenged the ability of the pipeline to re-analyze a dataset of the literature (Fig. 4 D, E, and F), namely the sorghum dataset of Mccormick et al. (2017), where a different 3D acquisition method was used and for which we had access to a limited amount of metadata. The third test involved the use of synthetic data (Fig. 4 G, H, I, and J) in order to directly compare outputs to ‘ground truth’ data that are perfectly known (Lobet, 2017), using maize plants generated with the ADEL maize simulation model (Fournier & Andrieu, 1998).

**Figure 4.**
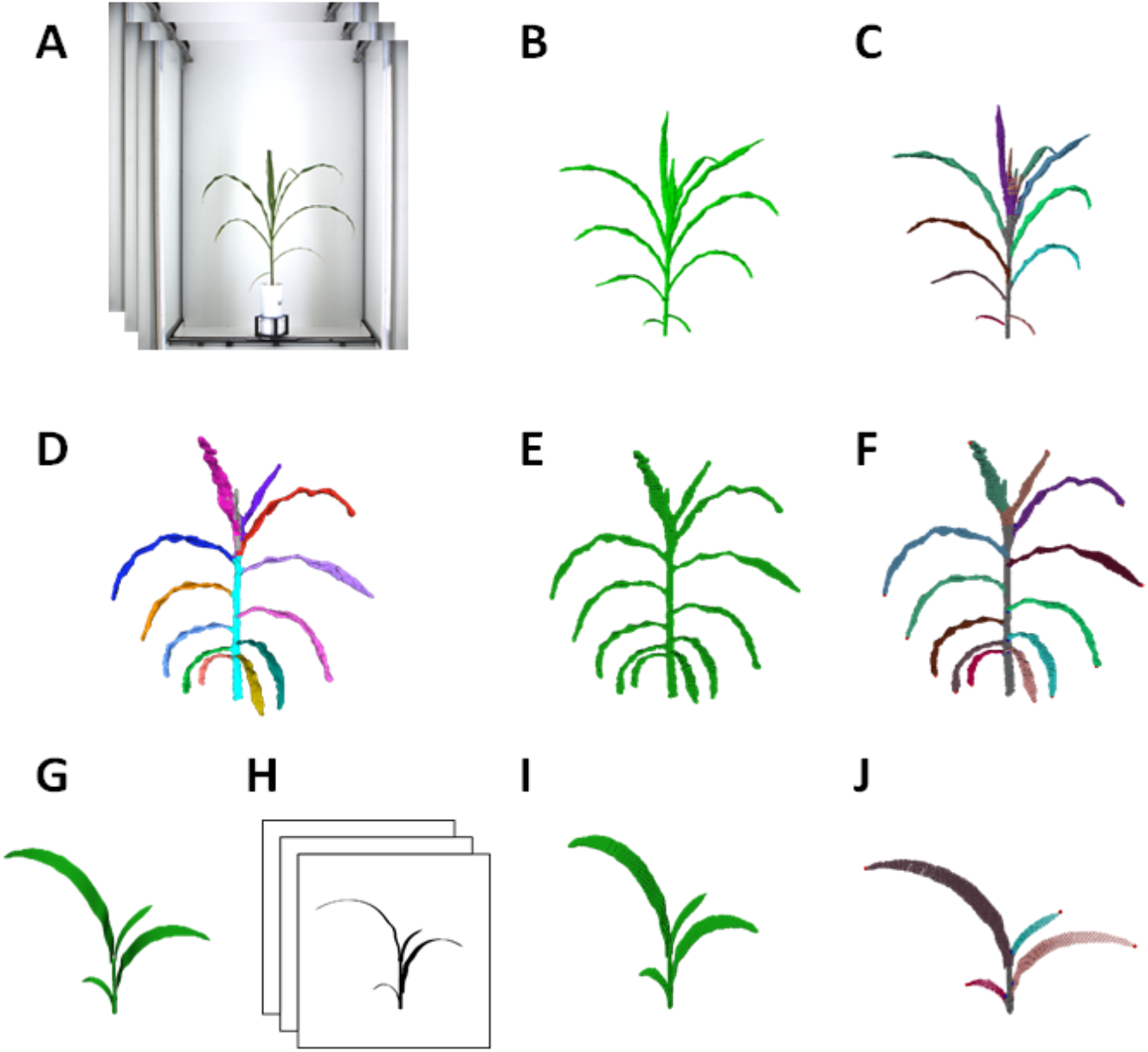
Snapshots of plants processed with Phenomenal for the three tests. Top : ground truth validation for a Phenoarch experiment where plants have been manually measured after imaging. (A) Input raw image from the platform (13 views), (B) 3D volume obtained with Phenomenal robust multi-view reconstruction (C) segmented plants. Middle: Re-analysis of a sorghum plant semi-automatically segmented by McCormick et al. (2016). (D)The original 3D mesh plant segmented from McCormick et al., (2016), (E) Initial 3D volume from McCormick et al., (2016) (F) segmented plant using *Phenomenal*. Bottom: Comparison with synthetic data. (G) 3D maize mesh generated from the *ADEL* maize model (Fournier and Andrieu, 1999), (H) binary images generated with *Phenomenal* projection on virtual cameral mimicking the *Phenoarch* 13 view acquisition system, (I) 3D volume computed using Phenomenal multi-view reconstruction (J) automatic segmentation with *Phenomenal*.

#### Comparison with manual measurement

The organ segmentation workflow was run on 39,548 reconstructed 3D maize plants from the experiment ZA16 of the *PhenoArch* platform (Table 1, Fig. 4A, B, and C), with a failure rate of 0.07%. On this experiment, 199,487 mature leaves and 168,404 expanding leaves were automatically detected. On the full set of reconstructed mature leaves, only 4.6% of them were artifacts (1.4% with maximal leaf width 20% larger than the actual measured maximum, 134 mm, or smaller than the actual measured minimum, 6 mm, and another 3.2% having width/length ratio larger than 0.26 or smaller than 0.009). In contrast, 15.6% of the detected expanding leaves had unrealistic morphological characteristics. This result is consistent with the intrinsic increase of complexity of the reconstruction in the upper part of the plant where leaves are rolled within each other. Manual leaf counts performed during the same experiment lead to a theoretical expectation of 184,190 fully expanded leaves (to be compared to the 192,305 realistic mature leaves found by the workflow) and 163,705 expanding leaves (versus 142,132). These numbers indicate that up to 96 % of the leaves were correctly detected, and that up to 96 % of them were correctly classified as mature. These performances can be compared to the 2D algorithm run by Das Choudhury (2018) on maize skeletons reconstructed from one side view selected out of two, which detected 4,011 leaves out of the 4,443 actual leaves tagged on images (90%).

*Phenomenal* organ segmentation analysis workflow was further validated by comparing image-based extracted features (Fig. 2) with detailed manual measurements performed on 1,218 leaves (i.e. 781 mature and 437 growing leaves) harvested from 127 plants at one date. On this dataset, the number of leaves was detected without bias, with a mean absolute error of 1.03 leaf (Fig. 5). This error was a bit higher than for the other tests (0.4 leaf for sorghum data and 0.2 leaf for synthetic data), probably related to the fact that this dataset focus only on one particular stage of development, characterized by a high proportion of growing leaves (33% of total), hence increasing the probability of occurrence of artefact detection.

**Figure 5.**
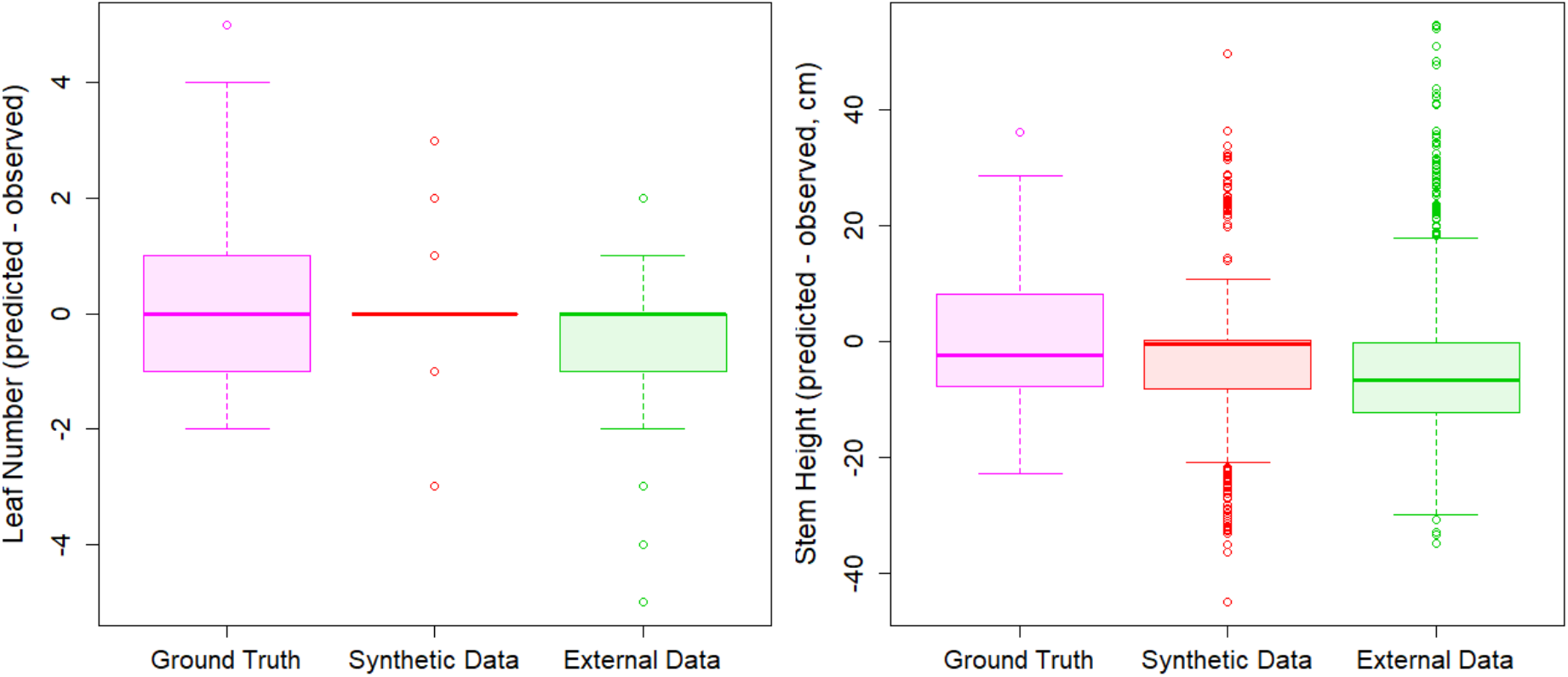
Boxplot of leaf number (left panel) and stem height (right panel) predictions for the 3 tested data set. (Pink) actual maize plants measured in the lab, (Red) synthetic maize plants generated with the ADEL, (green) segmented 3D sorghum provided by McCormick

As for leaf dimensions, amplitude and frequency of errors (Fig. 6 C, F and I) was smaller than previously reported error (McCormick et al. 2016) with a RMSD of 56 cm² for the area of 781 leaves (Fig. 6I, compared to a RMSD of 74.32 cm² for 140 leaves on McCormick study), leaf length (10.96 cm versus 7.94 cm, Fig. 6C) and leaf width (0.54 cm versus 1.84 cm, Fig. 6F). Part of this noise may originate from differences between the information present on the image and on the plant when it is measured in the lab. The tips of maize leaves, for instance, can often break when leaves are manipulated. This is consistent with leaf length being noisier than leaf area. Growing leaves were less accurately measured than mature ones, especially the shortest ones (Fig. S14 C). These leaves are those located in the center of the whorl, with a tip that cannot always be identified on silhouette images. The same occurred for leaf width (Fig.S14 F), and therefore leaf area (Fig. S14 I).

**Figure 6.**
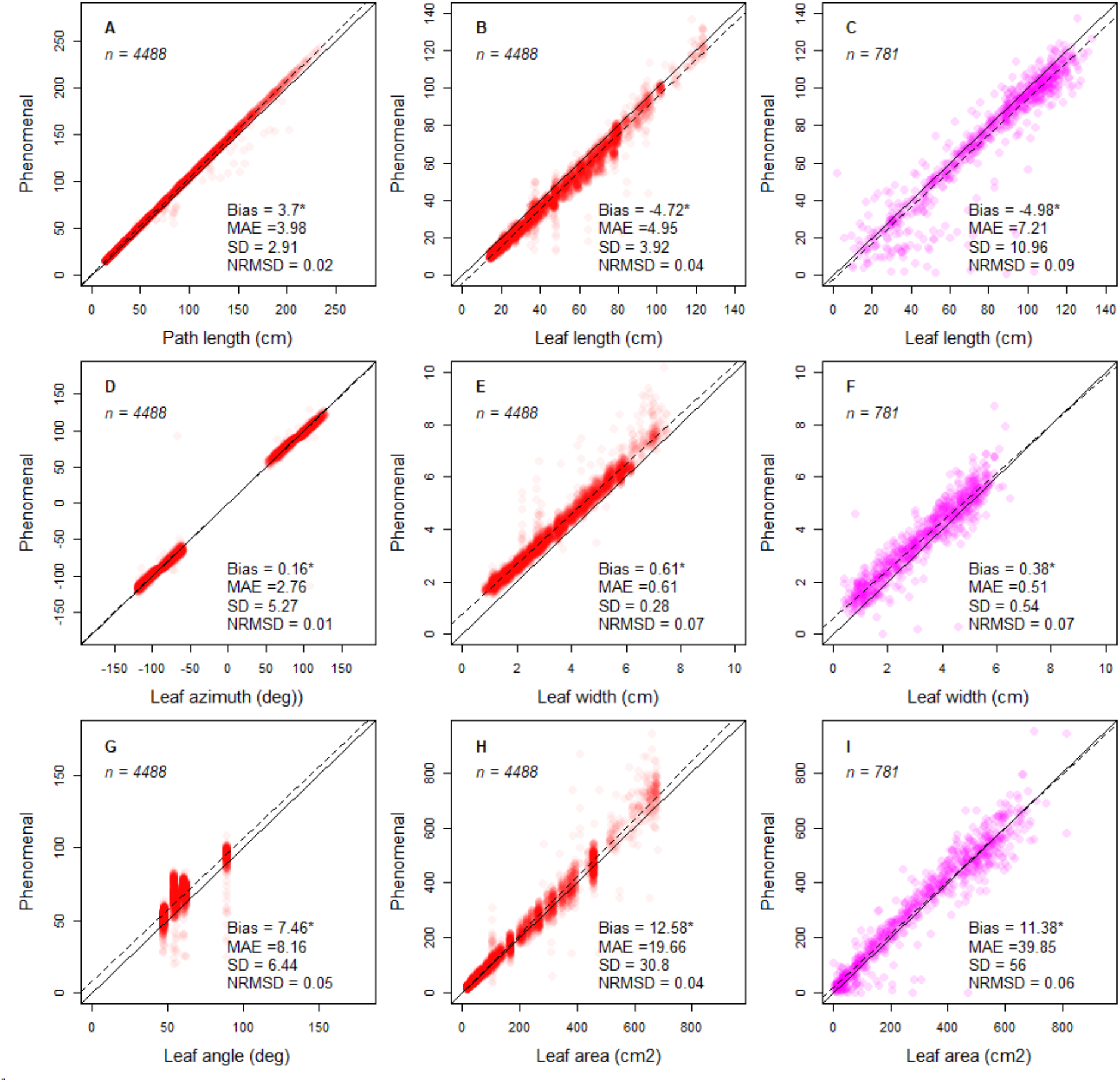
Comparison of image-based measurements with reference measurements for mature leaves detected by the *Phenomenal* organ analysis workflow vs. measurements for synthetic leaves and measurements made with a LICOR leaf area meter. Measurements with *Phenomenal* were performed on the 3D reconstructed plants. Plots visualize the bias between *Phenomenal* measurements and the ground truth values of synthetic leaves for skeleton path length, leaf length, leaf azimuth, leaf width, leaf angle and leaf area (red symbols; A-B, D-E, G-H). A second series of Plots were used to visualize the bias of *Phenomenal* measurements made on 3D reconstructed plants in comparison to manual measurements performed with the LICOR leaf area meter (pink symbols; C,F,I). An alpha-transparency of 0.05 and 0.15 have been applied to the plotted markers for red and pink symbols respectively. MAE: mean absolute difference; SD: standard deviation NRMSD: normalized root mean square difference (normalization by the range of values shown on x-axis)

#### Reanalysis of an existing dataset on sorghum

The organ segmentation workflow was challenged on a public dataset of 3D sorghum plants containing 1187 plants analyzed by McCormick *et al.* (2016). In their experiment, McCormick *et al.* provided semi-automatically segmented plants into leaves and stems. The Phenomenal workflow has been applied to process all this dataset automatically (Fig. 4D, E and F). The stem and leaf detection algorithm was considered by McCormick et al. (2016) as the main limiting step to the complete automation of their workflow on sorghum plants, and the authors kept this step manual to preserve the performance of their pipeline. The number of leaves identified was exact in 65% of cases, and for 87 % of the remaining plants the difference was only 1 leaf, i.e. similar error level as manual measurements (Fig. 5). Stem length estimates for this dataset were slightly biased, with a mean absolute error of 10.14 cm (Fig. 5). This corresponds to approximately 1.7 times the mean distance between the insertion points of successive leaves, suggesting that the estimation of stem endpoint by *Phenomenal* may be located one leaf below or above the manually determined insertion point. It should be noted that, unlike McCormick et al (2016), the error in the location of stem endpoint does not impact the detection and counting of individual leaves. The comparison of stem length estimates with reference values yielded very consistent results among datasets, in terms of bias and error (Fig. 5). The mean absolute error was found within one leaf to leaf interval for actual plants (sorghum or maize). Note that McCormick et al (2016) observed a bias of 5.75 and a root mean square error of 6.67 between manual measurement on actual plants and manual measurement on image for stem length.

#### Comparison with a synthetic dataset

The workflow was finally challenged by comparing image-based extracted features with corresponding known references on thousand synthetic 3D maize plants (corresponding to 4488 mature and 4291 growing leaves) obtained with the ADEL 3D-maize simulation model (Fournier & Andrieu, 1998) available in OpenAlea (Fig. 4 G, H, I, and J). This allows for testing the method accuracy for leaf number and leaf type detection, leaf length, maximal leaf width, leaf area, leaf insertion height, leaf azimuth and mean leaf angle for individual leaves of plants (Fig. 6). The segmentation of the leaves of the synthetic dataset resulted in an absence of bias and a mean absolute error of 0.2 leaves (Fig. 5), i.e. is two times more accurate than on the sorghum dataset. The comparison of stem length estimates with reference values yielded also very consistent results, with less than an internode error on the location the last ligulated leaf insertion. Morphology and size on mature leaves was accurately measured when compared with their synthetic counterparts (Fig. 6). The leaf azimuth angle and leaf path length (from the base of the plant to the tip of the leaf) were estimated almost without error by *Phenomenal* (1 and 2 % NRMSD respectively), indicating that the computed skeleton was reliable (Fig. 6A and D). The accuracy decreases slightly for estimation of the visible leaf length (Fig. 6B), with a mean absolute error of 5 cm (NRMSD = 4%), which almost always leads to an underestimation of the reference length. This difference is mostly due to errors in the localization of the leaf insertion point rather than errors in the localization of leaf tips (**Supplementary material S2**). The relative accuracy was lower for leaf width than leaf length (NRMSD = 7%, Fig. 6E), but it still allowed a good estimate of visible leaf area (Fig. 6H). Leaf angle estimations were less accurate (Fig. 6G), probably because plants are reconstructed with discrete voxels, therefore leading to unsmooth path in the skeleton.

This indicates that the binarization - which is exact on synthetic data (no background, movements or tutor) (Fig. 4 G, H, I, and J) - is probably an important point of attention for improvement of accuracy.

On growing leaves, we observed a slight increase of skeleton error for synthetic data (Fig. S3.1 A and D), as the NRMSD for leaf azimuth and leaf path length reached 3 %. This is consistent with the higher frequency of leaf crossing and pairing mismatch in this region of the plant. Errors on leaf length remain small (Fig. S3.1), with a 1.5 cm increase of error compared to mature leaves. The situation is very different, even on synthetic data, for leaf width and leaf area (Fig. S3.1E and H), with a doubling of the NRMSD (13%) compared to mature leaves, almost always resulting in an overestimation. Results of the comparison with actual data confirm and amplify these trends. Leaf inclination angles were accurate and comparable to mature leaf angle accuracy for a subset of the growing leaves, but highly variable for another subset (Fig. S3.1G). This is consistent with the increased frequency of tortuous skeleton path in the whorl.

### Coupling 3D shoot reconstruction with eco-physiological models

*Phenomenal* has been integrated in the *OpenAlea* framework, a platform dedicated to the analysis, validation and development of functional-structural plant models. This allows one to construct computational experiments mixing image analysis, 3D reconstruction and modelling steps. By connecting the output results of *Phenomenal* 3D reconstruction analysis workflow with ecophysiological models available in *OpenAlea*, we can compute foliar distribution (LAI), light interception and radiation-use efficiency. For example, the semi-automated workflow presented in Cabrera-Bosquet, et al. (2016) has been fully automatized using *Phenomenal* under *OpenAlea* to estimate light interception of individual plants reconstructed from *PhenoArch* platform (Fig. 7). This workflow was also used by Chen et al. (2018) on 3 different *PhenoArch* experiments (ZA13, ZB13 and ZA16) to analyze competition between plants, and by Lacube et al. (2017) to analyze the relation between leaf width and light interception. This workflow combines both the 3D reconstruction of plants with *Phenomenal*, the reconstruction of the virtual canopy in the greenhouse using *PlantGL* (Pradal, Boudon et al, 2009), and a radiative transfer model, *Caribu* (Chelle et Andrieu 1998), available in *OpenAlea*.

**Figure 7.**
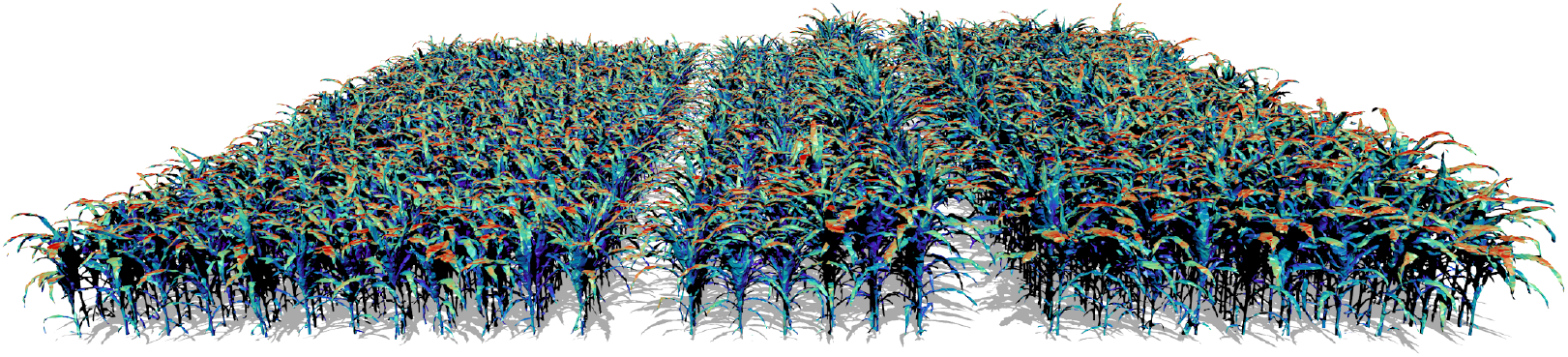
Example of extension of a *Phenomenal* pipeline with models available on the OpenAlea platform. The 3D maize canopy reconstruction pipeline built using *Phenomenal* was chained with the Caribu raycasting model available on OpenAlea to automate the simulation of light interception for the 1680 plants of the PhenoArch platform every day. The image shows the distribution of light irradiances (from blue to red) over a virtual field faithful to the actual arrangement of plants in the PhenoArch platform.

### A scientific workflow environment for HTP reproducible experiments

*Phenomenal* components, tools and algorithms are packaged as a Python package library under an open-source license using state-of-the-art computational science methodology. Developers can adapt the code and the design of analysis workflows to fit their experimental or processing needs. For quality assurance and continuous deployment, several tools have been set up. After each modification, a set of unit and functional tests are run on the cloud using continuous integration (https://travis-ci.org). When they succeed, the software is continuously deployed on the web for Linux, Windows and Mac operating systems on a *Conda* channel (Grüning et al., 2018). Documentation is also automatically rebuilt after each modification and updated on the web using the *readthedocs* service (https://readthedocs.org). To facilitate the installation of the software by end-users as well as administrators of computing infrastructures, *Conda* packages and Docker images are provided (Merkel, 2014). A set of tutorials are available as Jupyter Notebooks to enhance the computational understanding of the different steps by developers, the ability for end-user to reproduce the workflow, and the interaction with the 3D reconstruction through a web-based interface (Kluyver et al., 2016).

*Phenomenal* has been integrated into the OpenAlea SWfS, providing users with several major features. First, users are able to compose and interact with processing workflows within a visual programming environment: algorithms can be dragged and dropped in the user workspace, parameters can be modified and their behaviors interactively visualized (Fig. 1). Second, they have the ability to reproduce results (Cohen-Boulakia et al., 2017) obtained with the *Phenomenal* software which is guaranteed by the use of OpenAlea and its companion tools (Pradal et al., 2017). OpenAlea is indeed a SWfS able to keep track of the exact series of software used to design a scientific experiment and is provided with a provenance module, collecting parameter settings and datasets produced and processed during an execution. Moreover, to be able to process very large datasets obtained from phenotyping platforms, we have designed the infrastructure InfraPhenoGrid to distribute the computation of workflows on distributed computing facilities (Pradal et al., 2017; 2018; Heidsieck et al., 2019).

Finally, OpenAlea is built by an open-source community, which provides models and tools that are very valuable for plant phenotyping. The availability of ecophysiological models, as well as topological and geometrical data structures, makes it possible to implement a model-driven phenotyping approach. Moreover, the availability of several 3D plant architectural models allows to reuse the knowledge source for generating synthetic data for validation purposes on a wide range of condition and parameters.

## Discussion

In line with some other initiatives, such as *PlantCV* (Gehan et al., 2017) or *Scikit-image* (van der Walt et al., 2014), *Phenomenal* is a step towards an open-source ecosystem around high throughput image analysis of phenotyping platforms. However, to our knowledge, this is the first end-to-end open-source library that is able to analyze and reconstruct 3D models from images on large datasets in a reproducible way. While some algorithms are already available in other software, most of them have been enhanced to process actual HTP data automatically. RGB multi view acquisition systems like *PhenoArch* are widely spread, which makes our workflows probably also valid for processing data from other platforms. Whilst this is true for the image processing workflow itself, which can be run on every environment running Python, the 3D reconstruction workflow requires some adaptation due to specific acquisition and setup and data management strategies of each platform. In particular, the link with data management strategies is a key element to reach full automation, by retrieving camera calibration data and plant images as well as storing the resulted images and intermediate data. The complete integration of *Phenomenal* workflows in Python eases this automation.

Based on segmentation of 3D reconstruction, *Phenomenal* provides new traits for HTP experiments. First, it provides leaf counts on complex adult morphology, which is a key feature of phenological development and also important in the assessment of developmental stages (Erickson and Michelini, 1957; Arvidsson et al., 2011; Meicenheimer, 2014).

Second, it quantifies the total size and extents of the plant organs in the different dimensions. In graminea, like maize, being able to separate overall plant area growth from elongation growth is important, as elongation and widening are under different environmental controls and genetic regulations (Lacube et al., 2017). Although we do not assess it quantitatively, it was quite clear from our reconstruction that leaf thickness was clearly overestimated on the 3D reconstructed plants. This limitation remains even when refining the voxel size, and may therefore be intrinsic to the space carving method. *Phenomenal* also allows to quantify leaf angle distribution and its dynamics, which are key factors modulating light interception (Liu et al, 2017). These traits have been extracted by Lacube et al., 2017 but not in an automatic way, therefore limiting their use in large datasets. More originally, thanks to its connection to *OpenAlea* platform, *Phenomenal* allows retrieving some functional traits with models. Coupling with FSPM models could also improve the precision of analysis like Alvarez Prado et al. (2017) that inverted a simple evaporation model to get stomatal conductance estimates of thousands of plants in *PhenoArch*. Moreover, on maize plants, *Phenomenal* allows to separate the growing and mature leaves, and therefore allows the quantification of the whorl development. This structure is poorly characterized in the literature (Ruget et al., 1996), yet being involved in the regulation of the organ dimension, and is also the missing link between elongation and LAI models, as it determines the fraction of leaves exposed to light.

The extraction of biological traits is by nature more specific than neutral generic image analysis features such as object size or envelopes. For example, in the case of maize plants, it is important to separate mature from growing leaves to interpret the meaning of the leaf insertion height for correct architecture analysis. Leaf insertion visibility point is either at the base of the leaf or at the point of emergence. Such particularity makes HTP workflows less generic than pure image processing but provides more interpretable traits. However, while the segmentation has been developed for maize, it can help analyzing other cereal species that have a relative similar architecture (e.g. sorghum, wheat, barley, rice). For example, we have successfully tested the ability of the *Phenomenal* maize analysis workflow to process existing published dataset of sorghum from McCormick et al., (2016).

*Phenomenal* algorithms currently focuses on the independent analysis of images acquired through time on phenotyping platform. This allows to analyze kinetics for plant-level characteristics (e.g. volume, total length leaf counts). For organ-based traits, in which one wants to analyze for example the growth of individual leaves over time, a temporal analysis would require tracking. Tracking will also allow to get more robust analysis as some errors (e.g. in skeletonization) can be compensated over time. However, tracking of dynamic and deformable leaves is complex and provides further essential challenges, especially if robustness and reliability are key requirements, as is the case in fully automated assessments.

In this study, we use a virtual plant model only in the frame of the validation procedure. The generated plants and images could also be used to train deep learning networks (CNN) and retrieve more complex traits to overcome the current limitations of computer vision algorithms, mainly for tracking organ development (Pound et al, 2017; Ubbens et al, 2018). Reciprocally, linking models with *Phenomenal* can greatly improve models. FSPM have been of limited use, due to cost of parameterization/acquisition of data (Landl et al., 2018). HTP platforms inverse the problematic and open the way to calibrate FSPM for different species and/or different genotypes. This may allow predicting competition between plants and therefore find rules/models for community of plants. This is a current important limitation of crop models, which if addressed in term might then also help to further optimize holistic functional agro-ecological and environmental models on a landscape or field scale.

## Conclusions

*Phenomenal* provides a robust, end-to-end and open-source software library that allows setting up fully automated and data-intensive imaging analysis workflows for high throughput phenotyping platforms. It is able to manage large-scale data sets using distributed computing infrastructure. Based on botanical analysis of 3D reconstruction, *Phenomenal* provides new traits for assessment and analysis in HTP platforms. Accuracy on trait estimation is satisfactory for mature leaves but will require improvements for analysis of growing leaves. More originally, thanks to its connection to OpenAlea platform, *Phenomenal* allows to use models to estimate light interception and thus take into account the competition for light in analysis of platform experiments. *Phenomenal* algorithms currently focus on independent analysis of images acquired through time on phenotyping platforms. An important future evolution of *Phenomenal* is the development of tracking algorithms from sequences of images to enhance segmentation, 3D-skeletonization and trait extraction on various types of plants and take benefit of the redundancy of information to improve the accuracy of the analyses.

## Materials & Methods

### PhenoArch experiments and image acquisition system

Data from eight experiments conducted in the *PhenoArch* HTP platform (https://www6.montpellier.inra.fr/lepse/M3P, Cabrera-Bosquet et al., 2016) between summer 2012 and summer 2017 were used to test our workflow (Table 1). In all experiments, plants were grown in polyvinyl chloride (PVC) 9 L pots (0.19 m diameter, 0.4 m high) filled with a 30:70 (v/v) mixture of a clay and organic compost. Detailed descriptions of most of the experiments can be found in the literature (MA14: Lopez et al., 2015; ZB12, ZA13, ZB13 and ZA16 (Alvarez-Prado et al., 2017); ZA17 (Brichet et al. 2017); ZB14 and GA16, unpublished). The *PhenoArch* image acquisition system takes RGB color images (2056 x 2454 pixels) from the top and the side of the plant (Cabrera-Bosquet et al. 2016). By rotation of the plant, multiple images representing different lateral views are acquired (Fig. S1.1). The camera is precise enough for picturing the cabin without considerable spherical aberration (<0.1%, data not shown). Plants are usually imaged daily during nighttime, with customized number of images per plant. Typically, one image from top view and 2 to 24 images from side views are used for a given experiment.

### Multiple-camera calibration

Reconstruction of the 3D plant geometry from images requires the calibration of the imaging system. Although this procedure is standard in computer vision, we did not find a library that completely fit the specific requirements of phenotyping platforms (several cameras pointing at a well-controlled turning object). Therefore, *Phenomenal* library includes a specific calibration procedure, based on a generic optimization algorithm (Salvi et al., 2002), that allow retrieving at once the rotation angle, the axis of rotation and the intrinsic and extrinsic parameters of one or several cameras positioned around a rotation axis (**Supplementary Material S1-Camera Calibration**). These parameters allow to define the projection functions for the different views, and thus determine the pixel coordinate on any image of any point cabin. In the case of the *PhenoArch,.* The calibration was performed using images of a chessboard target taken every 3° from a side and top camera.

### Background subtraction

The details of the procedure we used to process *PhenoArch* images can be found in Brichet et al. (2017), and are summarized in **Supplementary material S1-Background subtraction**. Briefly, for side images, the segmentation combines a mean-shift threshold algorithm (Comaniciu and Meer, 2002) to a HSV threshold algorithm (Sural et al., 2002). For top images, we used a decision tree learning method (Breiman et al., 1984). In both cases, a median blur filter (Huang et al, 1979) was used for denoizing the result.

### Robust multi-view reconstruction

Based on camera calibration, the 3D plant architecture is reconstructed using a space carving algorithm (Kutulakos et al., 2000). The space carving algorithm detects the photo-consistent voxels in the scene (i.e. voxels for which all projections on binary images contain foreground pixels) and produces a 3D plant volume. To account for small deformations of plants induced by rotation of the pot during acquisition and errors in foreground / background segmentation, two adaptations have been made to this algorithm (**Supplementary Material S1-Robust multi-view reconstruction**). A first adaptation introduces a tolerance for non consistent projections on a given number of views. The second adaptation is to extend the 3D plant volume so that its re-projection along a particular direction perfectly match the observation.

The 3D surface is thereby computed from the volume with a marching cube algorithm (Lorensen & Cline, 1987) implemented in the VTK library (Schroeder et al., 2004). This allows to derive geometric traits such as surface area directly from this 3D representation. The size of the mesh surface is reduced using a decimation algorithm (Schroeder et al., 1992) and smoothed (Taubin et al., 1996) to simplify and improve further post-processing.

### 3D skeletonization

The segmentation of the 3D volume into individual organs is based the analysis of its skeleton, that is a one-dimensional graph that allows to capture the main topological relationships between different regions of the 3D volume (Tagliasacchi et al., 2013; Liu et al., 2011). Because standard skeletonization algorithms produce results that are too noisy, a new algorithm was designed in Phenomenal (**Supplementary Material S1 - 3D skeletonization**). It is based on an iterative selection of paths joining plant base to leaf tips, followed by a removal of voxels associated to this path. The result was further filtered to eliminate the paths associated to re-projections that always superposed to re-projection of other objects on the different views.

### 3D organ segmentation

The segments are finally partitioned into stem, mature leaves and growing leaves (**Supplementary Material S1 – Organ Segmentation**). The usual definition of stem is the morphological axe going from plant base to the position of the highest visible blade-sheath junction, named collar. As we do not directly detect collars on images, we defined the stem as the basal part of the path linking the base to the topmost point of the plant, up to the point where stem diameter suddenly and sharply increase due to the bunch of growing leaves forming the whorl. This definition is only a geometric proxy of the position of the highest collar, that works well during the vegetative stage.

### Deriving architectural traits at the organ level

Once the stem and individual leaves are segmented, morphological and architectural traits of these organs can be derived from their 3D-geometries. Traits extracted at whole plant level are *3D volume*, *maximal height*, *convex hull* and *projection area*.

The 3D volume is obtained with the number of voxels and their size. Maximal height of the plant is the Euclidean distance between the highest and the lowest voxels. Traits extracted from leaf levels are *leaf inclination angle* and leaf *azimuth* angle, *leaf length*, *maximum leaf width*, *average leaf width, leaf area* and *leaf volume* (Fig. 2). Azimuth is the angle between the mean vector of the leaf polyline and the *y* axis. Leaf inclination angle is the mean of angles between vectors of leaf polyline and z-axis. Length is defined as the sum of the Euclidean distances between two points along a polyline. The width is estimated along the leaf polyline by the maximal distance between two points in the set of voxels intercepted by a moving normal plane along the polyline. Leaf area is obtained by the integral of leaf width as a function of curvilinear abscissa of leaf polyline breakpoints. Traits extracted from the stem are *length*, *volume* and *diameter*. Length and volume are estimated similarly to leaves. To avoid artifacts due to stem enlargement at each leaf insertion (Brichet et al. 2018), the stem diameter is estimated only at positions of stem path where the moving normal plane pass through a local minimum (minimal peak of the diameter as a function of curvilinear abscissa curve).

### Validation with manual measurements

In order to compare manual and automatic measurements, 126 maize plants belonging to 26 genotypes with contrasting plant architectures were manually measured destructively at the end of experiment ZA16. For each plant, length, area, maximum and average width of the leaves and the stem were measured by a leaf area meter (LI-COR 3100 area meter, LI-COR, Lincoln, NE, USA). Before the destructive measurement, each plant was imaged with the *PhenoArch* acquisition system on 13 views (12 side views and 1 top view). From those images, the architectural traits were derived from the *Phenomenal* workflow configured with voxel diameter of 4 mm and a tolerance of photo-consistency equal to. To match the leaf data obtained from the workflow with the manual measurements, the visible leaf tips of a plant were first manually tagged on the best side view (the view with the binarized image with the largest convex hull). Then, the 3D leaf tips detected in the workflow were projected onto the same view. Finally, the distance between the projected tip and corresponding manually tagged tip were calculated (**Supplementary Material S2**).

To compare results from the *Phenomenal* workflow with manual measurements, comparisons were plotted (Fig. 6), which were optimized in regard for the visualization of large datasets using semi-transparent markers as proposed recently by Dhutia et al. (2017), with a chosen α-transparency value adjusted to the size and number of overlapping markers.

To allow to access both negative and positive errors, we also calculated the mean absolute error (MAE):

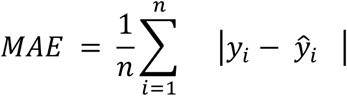

and the normalized root mean squared difference (nRMSD).

### Validation with synthetic data

The ADEL maize model (Fournier and Andrieu, 1999) simulates 3D geometry and topology of maize plants from a set of architectural parameters. The precise 3D geometry of each organ on the 3D mesh is therefore known, as well as the topological organization of the plant. We used the ADEL-maize model to simulate 1000 3D-maize architecture with random architectural parameters (Fig. 4 G, H, I and J). For each plant generated, model parameters were randomly and uniformly selected in specific intervals: plant height varied between 100 and 300 centimeters, the number of phytomers between 2 and 12 and base stem diameter between 1.25 and 7.5 cm. The other parameters are kept to ADEL default values. The geometry of these simulated plants were projected to different planes, which correspond to the ones of the *PhenoArch* phenotyping system, to create 2D binary images of these plants,. Then, the *Phenomenal* workflow was applied to those images to obtain 3D voxels, skeleton, organ segmentation and measurements of these plants. These results were compared to the synthetic data where everything is known (topology, geometry). This allowed us to validate the *Phenomenal* approach on a large set of conditions without expensive and complex manual data acquisition. Again to compare results from the *Phenomenal* workflow with measurements with synthetic data, comparisons were plotted as modified Bland-Altman plots as described in the previous paragraph, with a chosen α-transparency value adjusted to the size and number of overlapping markers.

## Acknowledgments

The authors acknowledge the support of France Grilles for providing computing resources on the French National Grid Infrastructure, of Institut Français de Bioinformatique project (IFB, PIA INBS 2012) for providing the cloud computing infrastructure, and the ‘Infrastructure Biologie Santé’ PHENOME-EMPHASIS project (ANR-11-INBS-0012) funded by the National Research Agency and the ‘Programme d’Investissements d’Avenir’ (PIA). MM has received the support of the EU in the framework of the Marie-Curie FP7 COFUND People Programme, through the award of an AgreenSkills fellowship under Grant agreement No. 267196. CP has received the support of the Thomas Jefferson Fund of the Embassy of France in the United States and the FACE Foundation. We are grateful to Claude Welcker for designing experiments and to all members at the *PhenoArch* platform for providing technical support, conducting the experiments and collecting data.

## Notes

This work was supported by the ANR-INBS projects PHENOME, EMPHASIS and IBC.

https://phenomenal.rtfd.io

